# Distribution of Epsilon-Polylysine Synthetases in Coryneform Bacteria Isolated from Cheese and Human Skin

**DOI:** 10.1101/2020.07.24.220772

**Authors:** Xinglin Jiang, Yulia Radko, Tetiana Gren, Emilia Palazzotto, Tue Sparholt Jørgensen, Tao Cheng, Mo Xian, Tilmann Weber, Sang Yup Lee

## Abstract

Epsilon-polylysine (ε-PL) is an antimicrobial commercially produced by *Streptomyces* fermentation and widely used in Asian countries for food preservation. Here we discovered a gene from cheese bacterium *Corynebacterium variabile* that showed high similarity to the ε-PL synthetase from *Streptomyces* in terms of enzymatic domain architecture and gene context. By cloning it into *Streptomyces coelicolor* with a *Streptomyces albulus* ε-PL synthetase promoter, we confirmed that its product is indeed ε-PL. A comprehensive sequence analysis suggests that ε-PL synthetases are widely spread among coryneform bacteria isolated from cheese and human skin; 14 out of 15 *Brevibacterium* isolates and 10 out of 12 *Corynebacterium* isolates contain Pls gene. This discovery raises the possibility that ε-PL as a bioactive secondary metabolite might be produced and play a role in the cheese and skin ecosystems.

**IMPORTANCE:** Every year, microbial contamination causes billions of tons of food wasted and millions of cases of foodborne illness. ε-PL is an excellent food preservative as it is potent, wide spectrum and is heat stable and biodegradable. It has not been accepted by all countries (e.g those in the EU) partially because it was not a natural composition of food but rather originated from the soil bacteria *Streptomyces*, a famous producer of various antibiotic drugs and toxins. The unexpected finding of ε-PL synthetases in cheese and skin bacteria suggests that ε-PL may naturally exist in cheese and on our skin.

---

Epsilon-polylysine (ε-PL) is a small cationic isopeptide made from the essential amino acid L-lysine (Fig. 1A). It exhibits antimicrobial activity against a wide spectrum of bacteria, yeast, and fungi and is heat stable and active in different food matrices (*1*). ε-PL has been a popular food preservative in Japan since the late 1980s, followed by Korea and China, and has been given Generally Regarded As Safe (GRAS) status in the USA. As a secondary metabolite, ε-PL was first discovered from the soil bacterium *Streptomyces albulus*, which is still used in its commercial production (*2*). Later, more producers of ε-PL were identified from the family of *Streptomycetaceae*, including the genera *Streptomyces* and *Kitasatospora*, and ergot fungi (*3, 4*). ε-PL is synthesized by a cell-membrane-bound nonribosomal peptide synthetase (NRPS)-like enzyme named ε-PL synthetase (Pls). The structure and mechanism of *Streptomyces* Pls has been well studied (*5*). Regulation of ε-PL biosynthesis and the natural role of this compound are less well understood.

**FIG. 1:**
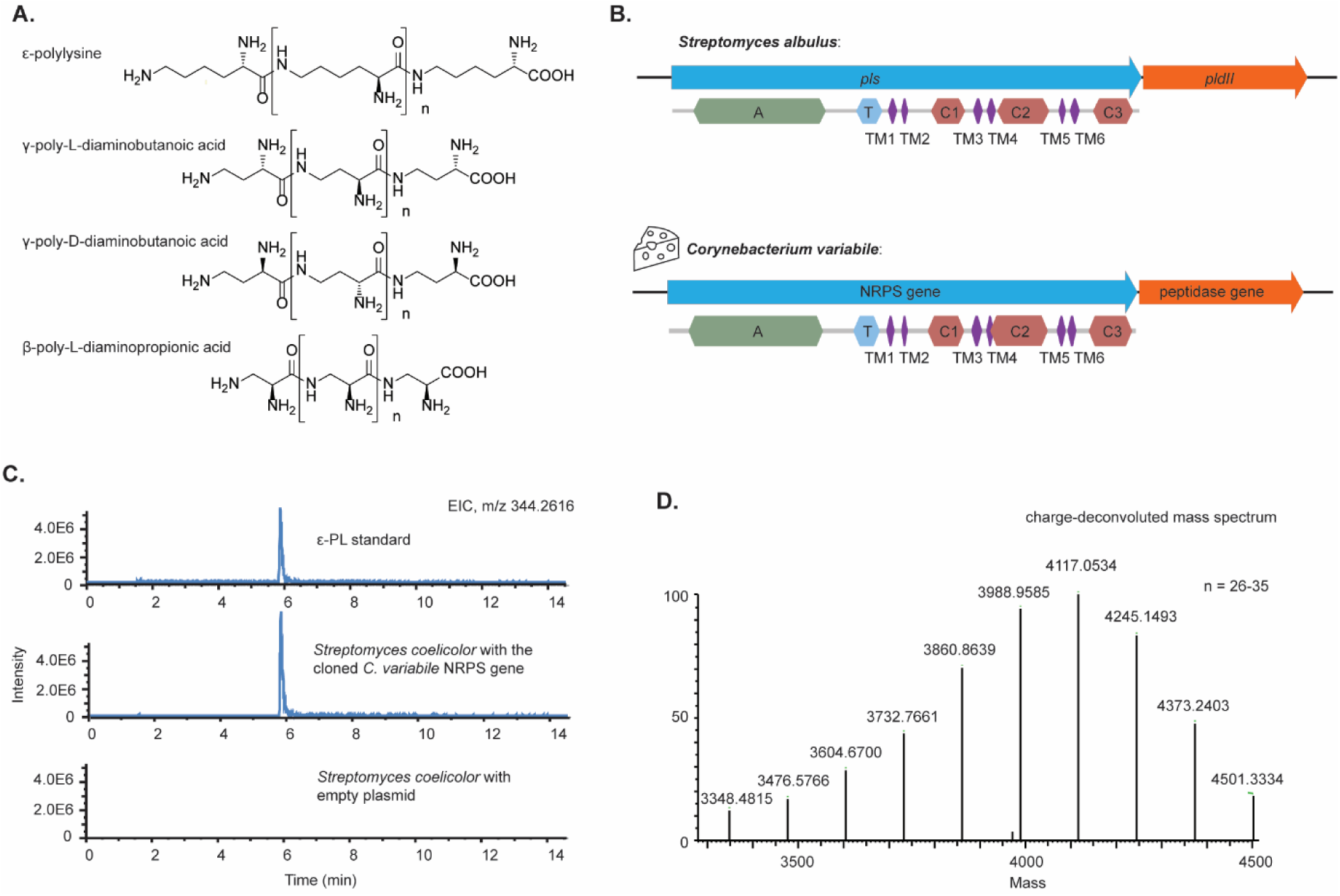
ε-PL synthetase gene from *Corynebacterium variabile*. (A) Chemical structures of ε-PL and related compounds. (B) Comparison between the ε-PL biosynthetic gene cluster from *Streptomyces albulus* and its homolog from *Corynebacterium variabile*. Arrows of the same colour indicate homologous genes. Domains of the synthetases were shown below the arrows. A, adenylation-domain; T, thiolation-domain; C1, C2 and C3, three C-terminal tandem domains related to acetyltransferases; TM1 to TM6, six transmembrane domains. Domain analysis was performed by InterProScan (*17*). (C) ε-PL was produced by *Streptomyces coelicolor* expressing the *C. variabile* NRPS gene. Bacterial culture extractions were analyzed by UHPLC-MS analysis. *S. coelicolor* with empty plasmid was used as negative control. (D) Precise MS of the product match with ε-PL of 26 to 35 residues.

Cheese prepared by fermentation of milk is an ancient food with a history of at least eight thousand years (*6*). The microorganisms on and in cheese and their secondary metabolites play key roles for the flavor, quality, preservation and safety of the cheese. When analyzing the genomes of cheese-isolated bacterium *Corynebacterium variabile* (*7, 8*) with antiSMASH (*9*), we noticed a gene encoding a NRPS-like enzyme with high similarity to *Streptomyces* Pls (table S1). InterProScan (*12*) results show that they share a unique domain architecture, which is not seen in typical NRPSs as found in the biosynthetic pathways of many peptide antibiotics, such as penicillin and vancomycin (*10*). The enzyme has a typical NRPS adenylation-(A)-domain for substrate activation and a thiolation-(T)/peptidyl carrier protein-domain for tethering the activated amino acid substrate. It, however, does not have the condensation domains (C-domain) nor thioesterase domains (TE-domain) of typical NRPSs. Instead, there are three tandem domains (C1, C2 and C3) related to acetyltransferases and six transmembrane (TM1 to TM6) domains separating the C1, C2 and C3 domains (Fig. 1B). Similar architectures can only be found in Pls related β-poly-L-diaminopropionic acid (β-PDAP) synthetase, γ-poly-L-diaminobutanoic acid (γ-PLDAB) synthetase and γ-poly-D-diaminobutanoic acid (γ-PDDAB) synthetase (*11-13*). β-PDAP, γ-PLDAB and γ-PDDAB are cationic isopeptides structurally similar to ε-PL (Fig. 1A). β-PDAP and γ-PLDAB are co-produced with ε-PL in different *Streptomyces* strains with higher antifungal activities and lower antibacterial activities than ε-PL (*11, 12, 14*). γ-PDDAB is produced by *Streptoalloteichus hindustanus* with strong antiviral activity and only weak antibacterial activities (*13, 15*). The *Corynebacterium* protein is more similar to Pls (sequence identity of 51%) than to the other three synthetases (32%, 33%, and 31%, respectively). Similar to the *Streptomyces* Pls gene, the *Corynebacterium* gene forms an operon with a peptidase gene, which is different from the gene contexts of the other three synthetases (figure S1). The *Streptomyces* peptidase (PldII) was shown to be a ε-PL degrading enzyme and postulated to have a self-protection function (*16*).

We tested ε-PL production from *C. variabile* using a two-stage culture method which was efficient in finding *Streptomyces* producers (*18*). However, no ε-PL production was detected in the culture. We reasoned that the cheese bacteria may have different regulation of ε-PL biosynthesis to the soil bacteria *Streptomyces*. Therefore, we cloned the *C. variabile* gene onto a plasmid with a pBAD promoter. The recombinant plasmid was transferred into model organism *Corynebacterium glutamicum*. However, again ε-PL production could not be observed.

In *Streptomyces*, the promoter sequence is critical for ε-PL production. It has been demonstrated that expression of *pls* in the native host *S. albulus* with an altered promoter did not lead to ε-PL production, but the use of the original promoter resulted in ε-PL production even in a heterologous *Streptomyces* host (*19*). Inspired by this, we cloned the *C. variabile* gene under the control of the *S. albulus pls* promoter and transferred the plasmid into *Streptomyces coelicolor* M145 which does not have an endogenous *pls* gene. Using this expression system, ε-PL production was confirmed by UHPLC and high-resolution mass spectrometry (Fig. 1C and D).

Next we investigated the distribution of *pls* in a genome collection of 156 bacteria isolated from cheese from Europe and United States (*20*). As the microorganisms from cheese are known to originate from or be related to animal and human skin microbiota (*21*) and ε-PL is also used in cosmetic products (*1*), we further included a genome collection of 124 microorganisms isolates from human skin (*22*). We used experimentally confirmed Pls protein sequences as query to do BlastP against the two collections with a cutoff of 40% sequence identity and 80% sequence coverage. Pls homologs were found to be concentrated in coryneform actinobacteria, including *Corynebacterium, Brevibacterium, Arthrobacter, Microbacterium, Glutamicibacter, Rhodococcus, Micrococcus* and *Dermacoccus*. No hit was found in bacteria from other phyla (Fig. 2A).

**FIG. 2:**
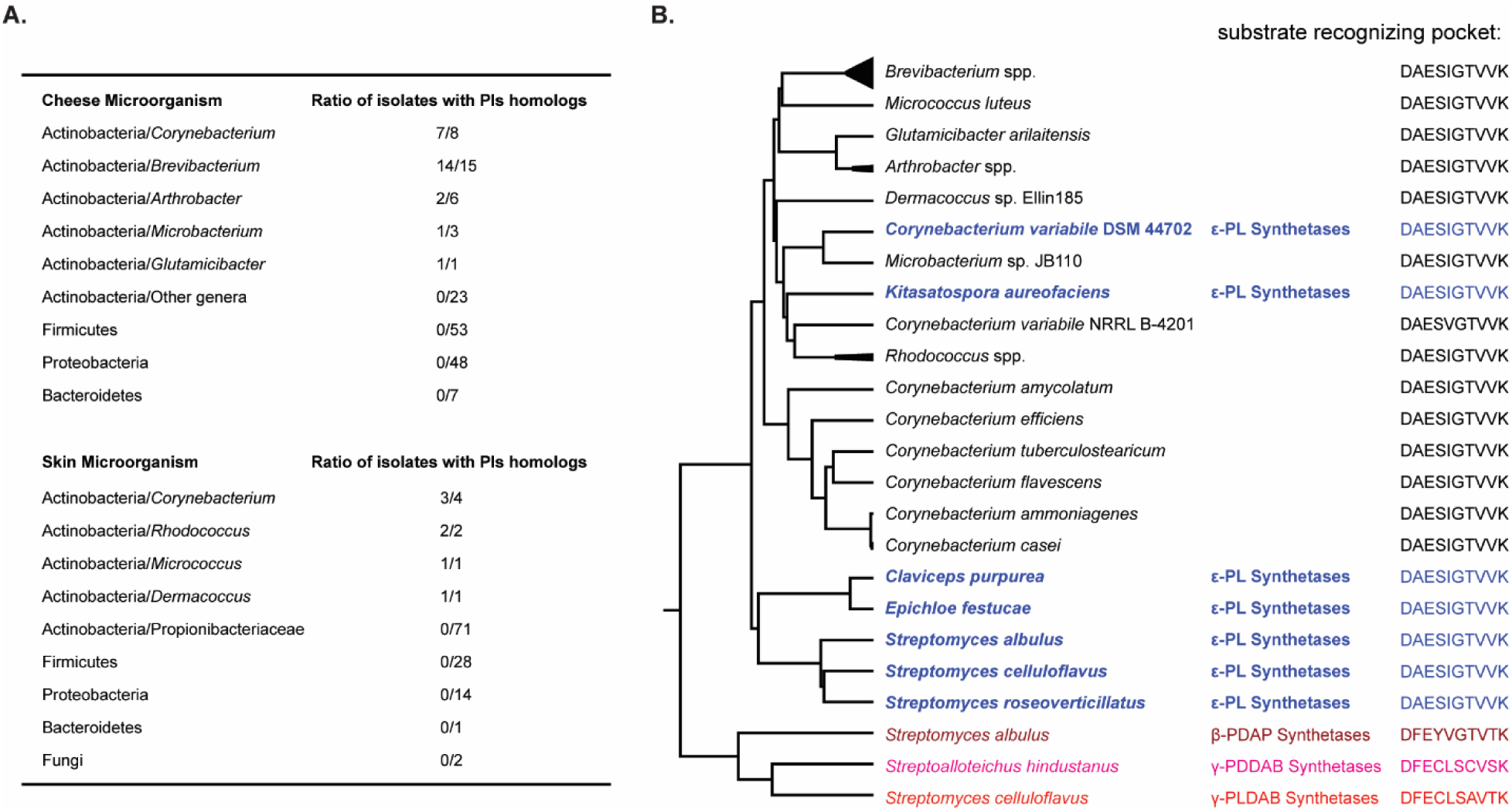
Pls distribution in microorganisms isolated from cheese and human skin. (A) Ratio of isolates with Pls homologs in different phyla and genera. (B) Protein phylogenetic tree and A-domain substrate recognizing pocket according to Stachelhaus et al (*23*). Experimentally confirmed Pls and related synthetases are shown in color. The homologs from cheese and skin bacteria were shown in black. Protein and genome access numbers, detailed blastP results and NRPSpredictor2 results are in Data set S1.

Phylogenetic analysis shows that all the coryneform bacteria proteins cluster together with the experimentally confirmed Pls from *C. variabile, Kitasatospora, Streptomyces* and fungi while the synthetases of the other three isopeptides are on more distant branches (Fig. 2B). Most importantly, NRPSpredictor2 (*24*) results show that the ten-residue substrate recognizing pocket (*23*) of the coryneform bacteria proteins are identical or highly similar to that of the confirmed Pls proteins but substantially different from that of the other three synthetases (Fig. 2B), which strongly suggests their substrate is lysine.

## Concluding remarks

In this study, we confirmed that cheese bacterium *C. variabile* has a ε-PL synthetase gene. We did not observe ε-PL production by *C. variabile* in our artificial culturing conditions. This is probably because the production of this secondary metabolite is regulated by a mechanism we are yet to understand, which is not uncommon for antimicrobial secondary metabolites. Furthermore, Pls were widely found in cheese- and skin-isolated coryneform bacteria. Most of the *Brevibacterium* and *Corynebacterium* isolates, which are among the most important microorganisms in cheese production, have Pls. It is possible that ε-PL naturally exists in cheese products and may have a role in cheese ecology, which requires further study.

## Bacteria

*Corynebacterium variabile* DSM 44702 was obtained from DSMZ. *Streptomyces coelicolor* M145 and *Corynebacterium glutamicum* MB 001 (DE3) were used as the heterologous hosts. *E.coli* DH5α was used for DNA cloning.

## Gene cloning

An expression vector pXJ0GC was developed from shuttle plasmid pAL374 (*25*) by adding a pBAD promoter and an apramycin selective marker. The *C. variabile* Pls gene was PCR amplified from *C. variabile* genomic DNA and cloned into pXJ0GC, resulting in plasmid pXJ146. pXJ146 was introduced into *C. glutamicum* by electroporation. A 450 bp DNA sequence containing the *pls* promoter from *S. albulus* was chemically synthesized and cloned together with the *C. variabile pls* into pRM4.3 resulting in plasmid pXJ155CV. pXJ155CV was introduced into *S. coelicolor* by conjugation. Primers DNA sequences and cloning details are summarized in Table S2.

## Culturing conditions

*Streptomyces* and *Corynebacterium* strains were maintained on ISP2 agar (BD Difco™). They were assayed for ε-PL production by a two-stage culture method (*18*). *S. coelicolor* strains were inoculated in M3G medium (*26*) at pH6.8 for 24 h at 30°C, then the pH was adjusted to 4.0 by HCl and cultured for another 3 days with shaking at 120 rpm. *Corynebacterium* strains were cultured similarly with GMPY medium (Malt extract 10 g/L; Peptone 10 g/L; Yeast extract 0.1 g/L autoclaved added with 10 g/L glucose as carbon source) and 1% arabinose was used for induction of gene expression in recombinant strains.

## Extraction

A Bond Elut LRC-CBA column (Agilent Part Number:12113037) was conditioned by washing with 5 ml methanol and then 5 ml water. Bacterial culture supernatant was adjusted to pH8 by NaOH and loaded on the column with a speed of 3 ml per min. The column was washed with 5 ml water and then eluted with 5 ml methanol twice. The elution was dried in a rotary evaporator at 38oC, redissolved and collected with 1 ml methanol and then concentrated to 50 μl using a vacuum centrifuge.

## UHPLC-MS analysis

UHPLC-MS analysis of the extract was performed on a Dionex Ultimate 3000 ultra-high-performance liquid chromatography (UHPLC) system coupled to a high resolution Orbitrap Fusion mass spectrometer (ThermoFisher Scientific, Waltham, MA, USA) and a UV/Vis diode array detector (DAD). Separate ESI+ and ESI-experiments were carried out with MS scan range of 100-1,000 Da. Injections of 8 µL of each sample were separated using a Waters Cortecs T3 column, 150 × 2.1 mm i.d., 1.6 µm particle size at a temperature of 35.0 °C and 0.35 ml/min flow rate. Elution was performed with 0.1 % formic acid in water (Mobile phase A) and 0.1 % formic acid in acetonitrile (Mobile phase B) in a multistep program: 0% B for 2.5 min, a linear gradient started from 0% to 100% B in 15 min, 100% B for 2 min, 0% B for 2 min.

## Bioinformatics

Protein domain analysis was performed by InterProScan. Phylogenetic analysis was done by MEGA-X using Muscle for multiple sequence alignment and Poisson model for UPGMA tree building. NRPS A domain substrate prediction was by NRPSpredictor2. For the cheese microorganism genome dataset,156 genomes were downloaded directly from the Supplementary Data 1 file of reference (*20*) and the amino acid sequences were extracted from the genbank format files using CLCgenomics (v.20.0). In addition to this, 47 of the 156 strains have genomes deposited in NCBI some of which are resequenced. Genes of them were downloaded from NCBI and included in the analysis. For the human skin microorganism genome dataset from reference (*22*), 124 genomes were downloaded from the NIH Human Microbiome Project (https://www.hmpdacc.org/hmp/catalog/grid.php?dataset=genomic). We downloaded the following Pls proteins to use as a reference: BAG68864.1, BAH85292.1, WP_014008652.1, AZL89021.1, CCE28893.1, and BBU42014.1. We then used BLASTP (v. 2.6.0+) with the following parameter to identify putative Pls in the downloaded amino acid datasets: -evalue 0.000001. We then extracted hits with at least 40% identity and at least 80% coverage of the reference gene.

## ACKNOWLEDGMENTS

This work was funded by grants of the Novo Nordisk Foundation [NNF10CC1016517 to SYL, NNF16OC0021746 to TW]. We would like to thank Dr. Pep Charusanti and Simon Shaw for valuable discussions and proofreading of the manuscript.

